# Astrocytes in the mouse visual cortex reliably respond to visual stimulation

**DOI:** 10.1101/429985

**Authors:** Keita Sonoda, Teppei Matsui, Haruhiko Bito, Kenichi Ohki

## Abstract

Astrocytes are known to contact with a great number of synapses and may integrate sensory inputs. In the ferret primary visual cortex, astrocytes respond to a visual stimulus with a delay of several seconds with respect to the surrounding neurons. However, in the mouse visual cortex, it remains unclear whether astrocytes respond to visual stimulations. In this study, using dual-color simultaneous *in vivo* two-photon Ca^2+^ imaging of neurons and astrocytes in the awake mouse visual cortex, we examined the visual responsiveness of astrocytes and their precise response timing relative to the surrounding neurons. Neurons reliably responded to visual stimulations, whereas astrocytes often showed neuromodulator-mediated global activities, which largely masked small periodic activities. Administration of the selective α1-adrenergic receptor antagonist prazosin substantially reduced such global astrocytic activities without affecting the neuronal visual responses. In the presence of prazosin, astrocytes showed weak but consistent visual responses mostly at their somata. Cross-correlation analysis estimated that the astrocytic visual responses were delayed by approximately 5 s relative to the surrounding neuronal responses. In conclusion, our research demonstrated that astrocytes in the primary visual cortex of awake mice responded to visual stimuli with a delay of several seconds relative to the surrounding neurons, which may indicate the existence of a common mechanism of neuron–astrocyte communication across species.

**Highlights:**

- We performed dual-color *in vivo* two-photon Ca^2+^ imaging of neurons and astrocytes.
- α1-adrenoblocker prazosin substantially reduced global astrocytic activities.
- Astrocytes showed weak but reliable visual responses in the awake mouse visual cortex.
- Astrocytic visual responses were delayed by 5 s relative to the neuronal ones.

## Introduction

Astrocytes play pivotal roles in supporting and maintaining neuronal and synaptic functions [1]. A single astrocyte in the cortex occupies an exclusive territory and contacts synapses with over 100,000 connections in mice, and up to 2,000,000 in humans [2]. Therefore, it has been suggested that the activities of astrocytes potentially reflect numerous synaptic activities in processing sensory information [3].

In the primary visual cortex (V1) of ferrets, it was reported that astrocytes responded to visual stimulation with a delay of 3–4 s compared to the surrounding neurons [4]. In contrast, previous studies in the mouse visual cortex reported that passive visual stimulation did not cause reliable and reproducible astrocytic sensory responses [5,6,7]. However, in these studies, astrocytic sensory responses were tested either in an anesthetized condition, which could have altered normal astrocytic activity [8], or in the awake condition for which astrocytic activities could be dominated by global events mediated by neuromodulators [5]. In the latter cases, such global activities in the awake condition could have masked small but reliable astrocytic visual responses.

In this study, we expressed highly sensitive calcium indicators, GCaMP6f [9] and R-CaMP2 [10] in astrocytes and neurons, respectively, to observe simultaneously the astrocytic and neuronal visual responses in the V1 of awake mice. We found that neuromodulator-mediated global activities dominated in the astrocytic activities. Pharmacological suppression of the global astrocytic activities revealed small but reliable astrocytic visual responses at their somata. Simultaneously monitored neuronal activity confirmed that the pharmacological manipulation did not significantly affect the neuronal visual responses. The astrocytic visual responses were delayed by approximately 5 s relative to simultaneously monitored neuronal visual responses. Together, these results showed that astrocytes in the mouse V1 were responsive to visual stimulations.

## Materials and methods

### Animals

We used 7–10 week-old C57BL/6 male mice (SLC, Japan) for all the experiments. All mice were maintained in the animal facility at the University of Tokyo, which housed 2–3 mice per cage in a temperature-controlled animal room with a 12 h / 12 h light / dark cycle. All experimental procedures used in this study were approved by the Animal Welfare Committee of the University of Tokyo.

### Preparation for chronic imaging and virus injection

During surgical preparation for window implantation and virus injection, mice were anesthetized with isoflurane (0.7–1.5%) delivered via a small nose cone. Skin incision was made at the midline of the mouse head. The periosteum and muscles disturbing surgery were removed from the skull. A custom-made metal head plate was attached to the skull with dental cement (Sun Medical, Japan). A small craniotomy (a diameter of 4 mm) was made over the left V1 (the center coordinated to + 3 mm lateral, ± 0 mm anterior from lambda). We used AAV1-CaMKIIpro-R-CaMP2-WPRE (2.0 × 10^13^ genome copies per ml, [10]) to express R-CaMP2 in neurons, and AAV5-GfaABC1D-cyto-GCaMP6f (1.81 × 10^13^ genome copies per ml, purchased from the University of Pennsylvania Vector Core, [9]) to express cyto-GCaMP6f in astrocytes. Although we also used AAV8-GFAP-hM3D-mCherry (2.2 × 10^12^ genome copies per ml, purchased from the Vector Core at the University of North Carolina at Chapel Hill) to express hM3D in astrocytes in some experiments, we did not use clozapine-N-oxide, which drove the Designer Receptors Exclusively Activated by Designer Drug system, in this study. The solutions of these AAVs were mixed at a volume ratio of 400 nl: 200 nl: 0 nl (or 100 µl in some experiments) in the order listed above. A total of 600–700 nl of this mixture was injected by a glass pipette using a Nanoject III (Drummond Scientific, PA) at 1 nl/s near the center of the craniotomy in the left V1 at a depth of 400 µm. The dura was not removed. The craniotomy was sealed with a round glass coverslip. The optical window was covered with Dent Silicone-V (Shofu, Japan) to prevent physical damage. After surgery, the mice were taken back to the cages and maintained for 6–10 weeks until two-photon imaging.

### In vivo two-photon Ca^2+^ imaging

On the imaging day, each mouse was fixed to stereotactic apparatus for a brief period of isoflurane administration. After the head fixation, we waited for at least 20 minutes before imaging to allow mice to recover fully from anesthesia. In imaging, great care was taken to prevent the visual stimuli from leaking into the light path of the microscope. For instance, a conical metal head cover was attached around the optical window. Ca^2+^ activities of neurons and astrocytes were recorded using a two-photon microscope (FVMPE-RS, Olympus, Japan) with a 25× water immersion objective lens (XLPLN25XWMP2, numerical aperture = 1.05, Olympus, Japan) at two excitation wavelengths of 940 nm for GCaMP6f and 1040 nm for R-CaMP2 (InSight DS Dual-OL, Spectra Physics, CA). The excited fluorescence was segregated with a 570-nm dichroic mirror and passed through the emission band-pass filters of 495–540 nm for GCaMP and 575–645 nm for R-CaMP. Note that the fluorescence signal from hM3D-mCherry, which was also expressed in astrocytes in some experiments, was too weak to disturb the recording of Ca^2+^ activities. Images were acquired in the layer 2/3 (depths of 160–230 µm from the pia mater) of the left V1 at 256 × 256 pixels at 0.41 s per frame with a galvanometer scanner.

### Visual stimulation

Visual stimuli were presented on a 32-inch LCD display (ME32C, Samsung, Korea) using PsychoPy2 [11]. The center of the display was positioned at 18 cm from the right eye of each mouse. Square-wave drifting gratings were presented sequentially in 8 directions of motions with 45-degree steps (spatial frequency = 0.04 cycles per degree, temporal frequency = 2 Hz). These 8 patterns were presented for 4.1 s (10 frames) each, interspersed with gray blank stimuli for 4.1 s (10 frames) or 12.3 s (30 frames) as inter-stimulus intervals (ISI). A trial consisting of these stimuli was repeated 10 (ISI = 12.3 s) or 20 times (ISI = 4.1 s). The red light component was set to zero in order to minimize contamination of the fluorescence.

### Pharmacology

Prazosin hydrochloride (Tokyo Chemical Industry, Japan) was dissolved in saline, and a 0.1 mg/ml solution was prepared. This solution was administered at a dose of 3 mg/kg intraperitoneally > 15 minutes before imaging.

### Data analysis

All data were analyzed using custom-written programs in MatLab (Mathworks, MA). A series of R-CaMP images were registered with the algorithm using the ‘dftregistration’ function [12]. The calculated integer values of pixels of xy shifts in individual frames were also applied to register a series of GCaMP images. The registered images of 256 × 256 pixels were trimmed to 240 × 240 pixels. Raw timecourses in field-of-view (FOV) were obtained by averaging the signal intensity of all pixels in the images. Slow drift of the baseline signal over the course of minutes was removed using a low-cut Gaussian filter with cut-off periods of 400 frames for R-CaMP and 3200 frames for GCaMP. Then, the timecourses were smoothed with a 10^th^ order Butterworth high-cut filter with a cut-off period of 0.41 s. For the experiment where we pharmacologically blocked the effect of noradrenaline, we applied a different low-cut filter (cut-off period, 70 frames) to astrocytic FOV timecourses in order to detrend the signal more strongly. The trials with residual global trends in astrocytic FOV timecourses were discarded from the analysis by visual inspection. An F0 value was defined as the 30-percentile value of each smoothed timecourse. Timecourses were divided by F0 values and subtracted by 1 to obtain dF/F0. To make astrocytic dF/F0 images, the same processing mentioned above was applied to individual pixels without spatial averaging. Then, images at individual frames were extracted and presented as pseudo-colored maps. An astrocytic ‘event’ was defined as a time-window in which an astrocytic FOV timecourse exceed 5% for more than 20 consecutive frames. Pixel-based hue-lightness-saturation (HLS) orientation maps of neurons were obtained as described previously [13]. The visual responses of neuronal R-CaMP signals were determined by the mean dF/F0 values during the 10 frames in which drifting gratings were presented relative to the baseline, which was defined as the mean dF/F0 value of the last 5 frames of each blank period immediately before the presentation of drifting gratings.

To analyze the R-CaMP signal of individual neurons, circular regions of interest (ROI) with a diameter of 16 pixels (approximately 16 µm) were set manually by referring to the average images, the spatially high-passed images of FOV, and the HLS orientation maps. A ring-shaped ROI with a width of 2 pixels outside each circular ROI was regarded as a neuropil ROI. Raw neuronal and neuropil timecourses were obtained by averaging the signal intensity of all pixels in the neuronal and neuropil ROIs. De-noised neuronal timecourses were then obtained by subtracting from the raw neuronal time courses the neuropil timecourses multiplied by 0.3 [13]. Then, the same procedures as we used for FOV timecourses (low-cutting, high-cutting, and defining F0) were applied to obtain the timecourses of dF/F0 for individual neuronal ROIs.

To analyze the GCaMP signal in the astrocytes, for each scan, we first made a pixel-based map of standard deviation across time. These standard deviation maps were smoothed with median filters (10 × 10 pixels, 20 × 20 pixels, 40 × 40 pixels) repetitively, and thresholded by 1.5. Contiguous regions above the threshold with the size larger than 3 pixels were regarded as astrocytic ROIs. Astrocytic ROIs for somata and processes were determined by visual inspection. Astrocytic timecourses for individual ROIs were obtained by averaging the signal intensity of all pixels in the ROIs. The same procedures used for the astrocytic FOV timecourses (low-cutting, high-cutting, and defining F0) were applied to obtain the timecourses of dF/F0 for individual astrocytic ROIs.

### Statistical analysis

Paired *t* tests were used to compare the datasets before and after prazosin administration. The sample size was defined as the number of FOVs. Throughout the study, p < 0.01 was considered statistically significant.

## Results

Using dual-color simultaneous *in vivo* two-photon Ca^2+^ imaging, we monitored the activities of R-CaMP-expressing neurons and GCaMP-expressing astrocytes in the V1 of awake mice (Fig. 1A). Visual stimulations with drifting gratings reliably evoked neuronal responses (Fig. 1B, top, a timecourse of all trials; Fig. 1C, top, trial-averaged timecourses), whereas astrocytic activities were dominated by global activities that did not resemble the simultaneously recorded neuronal activity (Fig. 1B, bottom, a timecourse of all trials; Fig. 1C, bottom, trial-averaged timecourses). Although small periodic astrocytic activities were visible (Fig. 1B, bottom, inset), the presence of the global astrocytic activities often masked such small periodic astrocytic activities. These global astrocytic activities are known to be evoked by noradrenaline released by projection fibers from the Locus Coeruleus [14] and are blocked by administration of the selective α 1-adrenergic receptor antagonist prazosin [15]. Therefore, we used prazosin administration to prevent global astrocytic activities from masking the small periodic astrocytic activities.

**Fig. 1.**
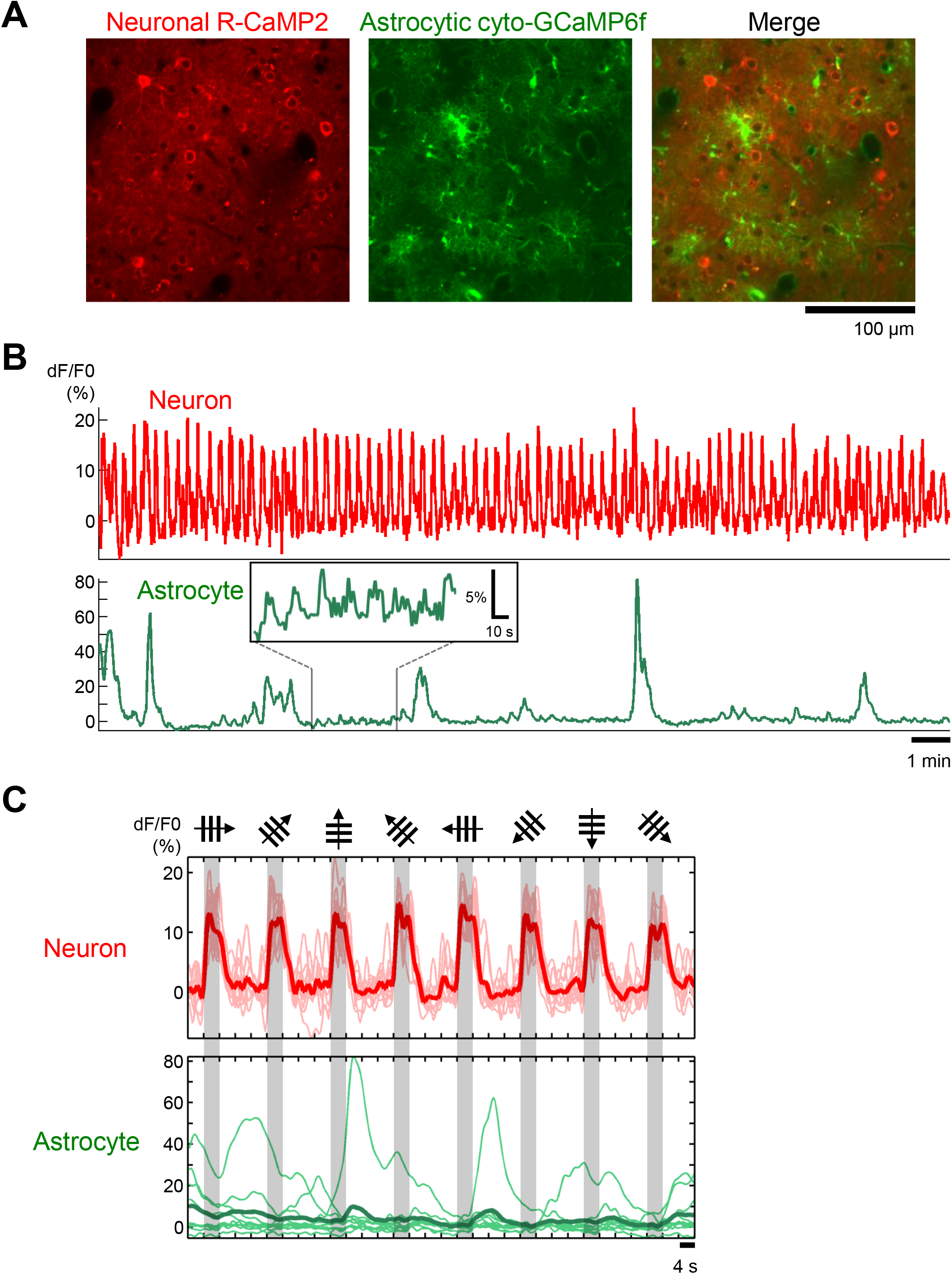
Simultaneously monitored neuronal and astrocytic activities in the awake mouse visual cortex. (A) Average images of R-CaMP2-expressing neurons (left), cyto-GCaMP6f-expressing astrocytes (middle), and both images merged (right). (B) Timecourses of the FOV in (A). Top, neuron (red). Bottom, astrocyte (green). Inset, a magnified trace. (C) Trial-averaged timecourses (thick traces) of the neuronal R-CaMP2 signal (top, red) and the astrocytic GCaMP6f signal (bottom, green) in the FOV in (A). Thin traces, individual trials. Gray shades, visual stimulation periods. Drifting directions of the gratings are shown on top of the gray shades.

As expected, the global activities of astrocytes were substantially reduced after administration of prazosin (Fig. 2A). Quantification of event frequencies revealed a statistically significant reduction of global astrocytic activities by prazosin administration (Fig. 2B; pre, 0.41 ± 0.05 events/min; post, 0.06 ± 0.02 events/min; mean ± standard error of the mean (s.e.m.), p < 0.0001, paired *t* test, n = 13 FOVs, 9 mice). Next, we examined whether the visual responses of neurons were preserved in the presence of prazosin. Both orientation maps and individual neuronal timecourses were similar before and after prazosin administration (Fig. 2C,D). The neuronal visual responses were not markedly altered by prazosin administration (Fig. 2E; pre, 4.60 ± 0.82%; post, 3.79 ± 0.76%; mean ± s.e.m., p = 0.02, paired *t* test, n = 13 FOVs, 9 mice). Thus, we verified that prazosin administration did not significantly alter the neuronal visual responses while abolishing the global astrocytic activities.

**Fig. 2.**
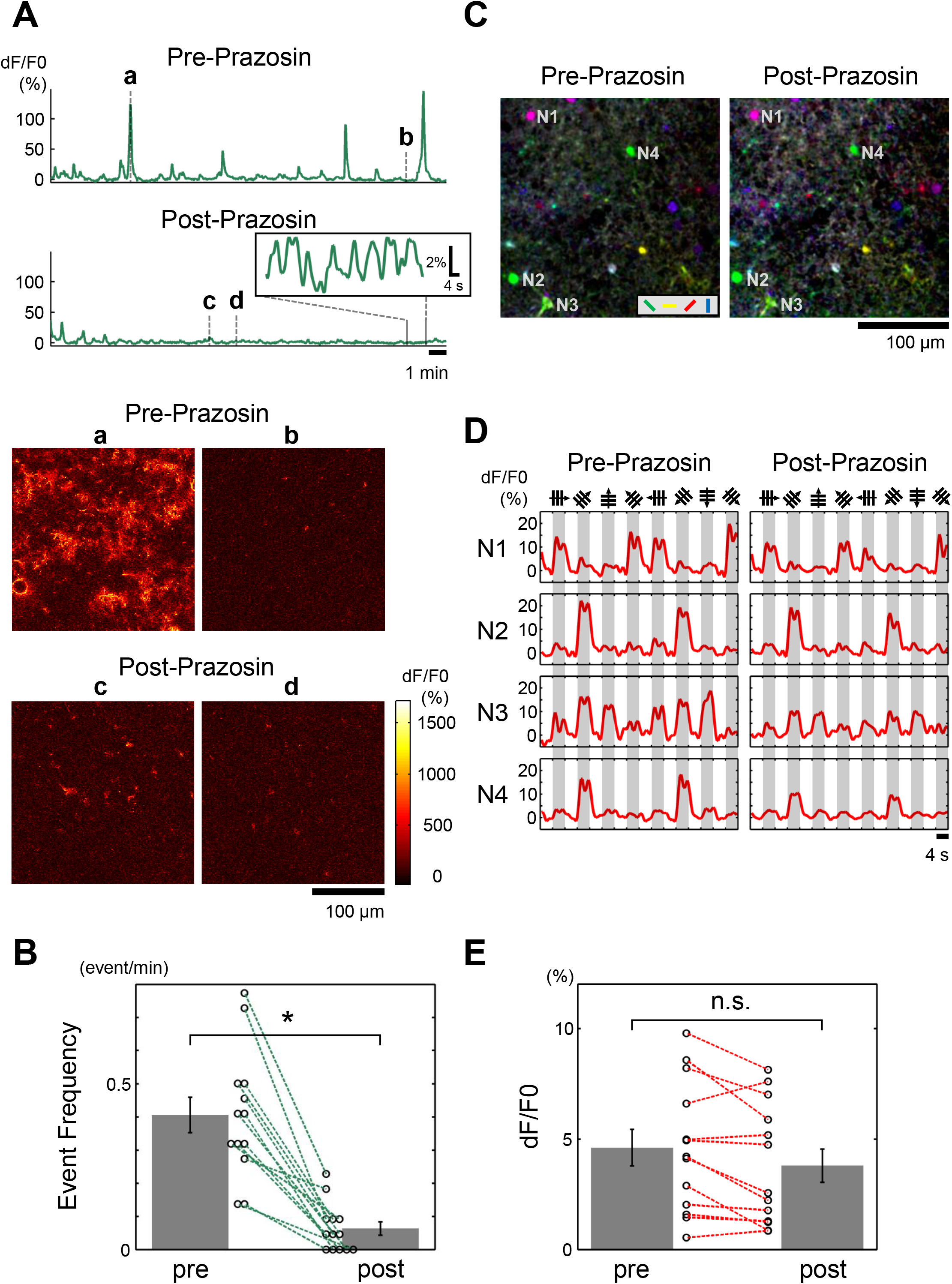
Effects of α1-adrenoblocker, prazosin on astrocytic and neuronal activities. (A) Timecourses of astrocytic GCaMP6f in the same FOV before (top) and after (middle) prazosin administration. Inset, magnified trace. Note that small periodic astrocytic activities are preserved after prazosin administration. Images at the bottom show dF/F0 images at representative time points (a–d). (B) Astrocytic event frequencies before (pre) and after (post) prazosin administration. Error bars, s.e.m. Circles, individual FOVs. *, p < 0.01 (paired *t* test). (C) Pixel-based HLS orientation maps of neurons before (left) and after (right) prazosin administration. Data from a different mouse as used in (A) and (B). Colored bars in the box indicate coloring of preferred orientation. (D) Trial-averaged timecourses of the individual neurons (N1 to N4) shown in (C), before (left) and after (right) prazosin administration. Gray shades, visual stimulation periods. Drifting directions of the gratings are shown on top of the gray shades. (E) Visual responses (dF/F0) of the neuronal R-CaMP signal before (pre) and after (post) prazosin administration. Error bars, s.e.m. Circles, individual FOVs. n.s., not significant.

In ferrets, it was reported that astrocytes in the V1 showed visual responses delayed by 3–4 s relative to the surrounding neurons [4]. Therefore, we sought to study the possible visual responses of astrocytes under prazosin administration in mice. We found that the astrocytes showed small but reliable visual responses, accompanied with some delays relative to the onset of visual stimulations, as in ferrets (Fig. 3A). Unlike neurons in the V1, these astrocytic visual responses were not selective for stimulus orientation. These astrocytic responses were mostly seen in the somata of the astrocytes (Fig. 4). In contrast to the somata, the processes showed activities thought to be unrelated to visual stimulations (Fig. 4). Taken together, prazosin administration unmasked the astrocytic visual responses, especially at their somata.

**Fig. 3.**
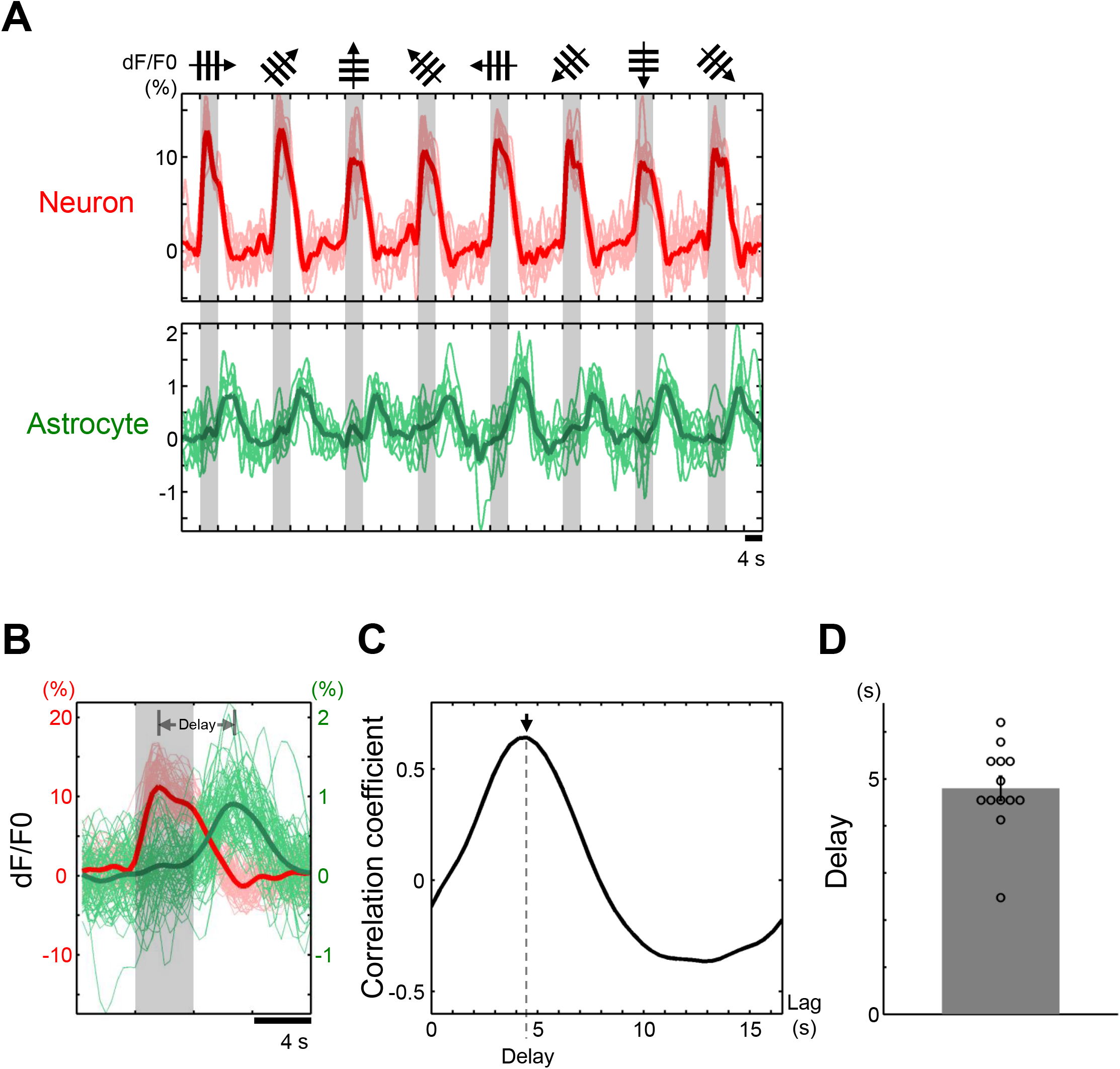
Astrocytes show delayed visual responses relative to neurons. (A) Timecourses of neuronal (top, red) and astrocytic signals (bottom, green) in the representative FOV under prazosin administration. Thin traces, individual trials. Gray shades, visual stimulation periods. Drifting directions of the gratings are shown on top of the gray shades. (B) Stimulus-averaged timecourses of neuronal (red) and astrocytic signals (green). The same data as in (A). (C) Cross-correlation analysis of neuronal and astrocytic signals in the same FOV. An arrow and a dotted line indicate a peak of correlation coefficient, i.e. a delay of visual responses in astrocytes relative to neurons. (D) Population data of the delay calculated with cross-correlation in 13 FOVs. Error bars, s.e.m. Circles, individual FOVs.

Finally, by taking advantage of the dual-color imaging, we investigated the precise temporal relationship between neuronal and astrocytic visual responses. Stimulus-averaged timecourses clearly showed a delay of several seconds between the neuronal and astrocytic visual responses (Fig. 3B). Cross-correlation analysis also revealed a delay of approximately 5 s (Fig. 3C). In the population data, the average delay was calculated to be 4.80 ± 0.25 s (Fig. 3D, mean ± s.e.m., n = 13 FOVs, 9 mice), comparable to the results of ferrets [4]. Taken together, it was indicated that astrocytes in the mouse V1 responded to visual stimuli with a delay of approximately 5 s relative to the surrounding neurons.

**Fig. 4.**
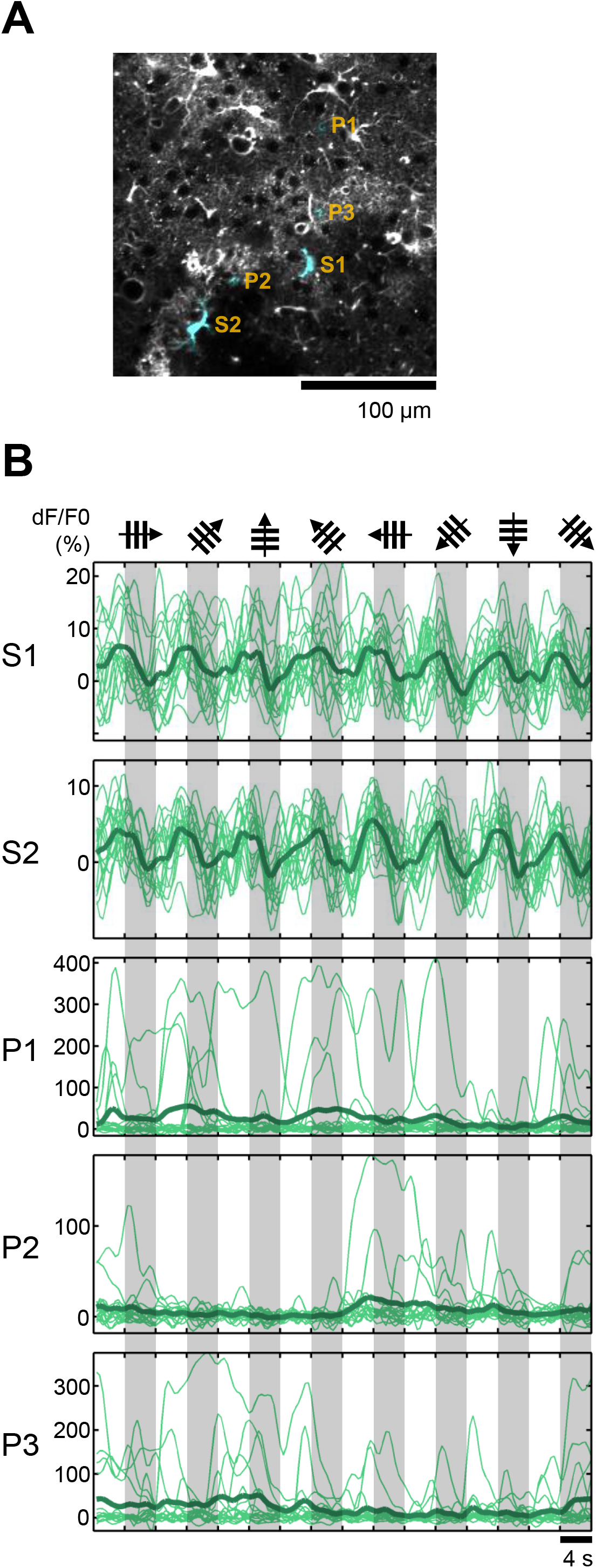
Astrocytes show visual responses at their somata. (A) Average images of cyto-GCaMP6f-expressing astrocytes. Regions colored in blue indicate the individual ROIs in astrocytes at the somata (S1 and S2) and processes (P1 to P3). (B) Timecourses of the ROIs shown in (A). Thin traces, individual trials. Gray shades, visual stimulation periods. Drifting directions of the gratings are shown on top of the gray shades.

## Discussion

In this study, using dual-color simultaneous *in vivo* two-photon Ca^2+^ imaging, we demonstrated the visual responses of astrocytes in the V1 of awake mice. These occurred mainly at astrocytic somata, and were delayed by approximately 5 s relative to the neuronal visual responses.

This is the first work to succeed at detecting the small but reliable visual responses of astrocytes. As in ferrets, the astrocytic visual responses were mainly found in the somata. In a previous study of mouse V1, Bonder et al. reported the absence of astrocytic visual responses in the somata [6]. The discrepancy may be attributed to their use of chlorprothixene as the sedative drug and isoflurane as the anesthesia, which might have altered the normal Ca^2+^ activities of astrocytes [8]. Previous studies in the awake mouse V1 also reported unreliable visual responses in the astrocytes [5,7]. The critical technical difference between these studies and the present study is that we used prazosin administration to suppress global astrocytic activities. The pharmacological treatment allowed us to isolate the small astrocytic visual responses from the neuromodulator-mediated global activities and precisely examine their timing relative to the visual stimulation and the neuronal responses.

Previous studies reported researching astrocytic Ca^2+^ activities in the mouse somatosensory cortex. In anesthetized mice, Wang et al. initially demonstrated that whisker stimulation evoked astrocytic Ca^2+^ activities broadly in the barrel cortex [16]. A subsequent study showed that such sensory evoked broad astrocytic Ca^2+^ activities were mediated by noradrenaline delivered from the Locus Coeruleus [14]. Similarly in the awake mice, the global activities of astrocytes caused by sensory stimulation were mediated by noradrenaline delivered from the Locus Coeruleus and were abolished by administration of prazosin [15]. Notably, they also reported that astrocytes did not respond directly to sensory stimulation under prazosin administration. In our study, the use of highly sensitive calcium indicator GCaMP6f likely allowed us to detect small but reliable astrocytic activities. Alternatively, neuron–astrocyte communication may be different in different sensory cortices [17].

Compared to ferrets, despite the similarity of timecourses, the amplitude of astrocytic visual responses were much smaller in mice. A potential explanation for this may be the difference of neocortical functional architecture in mice and ferrets [3]. The drifting gratings, which were used as the visual stimulus both in the present study and in the ferret study [4], evoked distinct spatial patterns of neuronal activities in the mouse and the ferret visual cortex. In the mouse, neuronal responses were sparse and had a salt-and-pepper arrangement. On the other hand, in the ferret, neuronal responses were dense and had a columnar arrangement. Therefore, a larger fraction of neurons surrounding an astrocyte is likely to be active upon visual stimulation in ferrets than in mice, which may be directly related to the difference in the amplitude of astrocytic visual responses. Future studies are needed to address this point to understand fully the species differences in neuron–astrocyte communication.

## Conflicts of interest

None.

## Acknowledgements

We thank Ayako Honda, Yumiko Sono, Mayumi Nakamichi, Ai Ohmori, Tomoko Inoue, Miki Saito and Aiko Hayashi for animal care, Takashi Kawashima, Kenta M Hagihara, Ayako Hayashi, Gen Ohtsuki, Kenji Hayashi and all members of Ohki laboratory for support and discussions. This work was supported by grants from Japan Society for the Promotion of Science (JSPS) KAKENHI (Grant number 25117004 and 25221001 to K.O.; 16H01328 and 17H06312 to H.B.; 17K14931 and 18H05116 to T.M.), Core Research for Evolutionary Science and Technology (CREST)—Japan Agency for Medical Research and Development (AMED) (to K.O.), Brain Mapping by Integrated Neurotechnologies for Disease Studies (Brain/MINDS)—AMED (to K.O.), Strategic International Research Cooperative Program (SICP)—AMED (to K.O.), Asashi Glass Foundation, Takeda Science Foundation, and Japan Foundation of Institute for Neuropsychiatry (to T.M.).

